# Identification of Periostin as a critical niche for myofibroblast dynamics and fibrosis during tendon healing

**DOI:** 10.1101/2023.07.21.550090

**Authors:** Jessica E. Ackerman, Emmanuela Adjei-Sowah, Antonion Korcari, Samantha N. Muscat, Anne E.C. Nichols, Mark R. Buckley, Alayna E. Loiselle

**Affiliations:** Center for Musculoskeletal Research, University of Rochester Medical Center, Rochester, NY; Current affiliation: NDORMS, University of Oxford, Oxford, United Kingdom; Department of Biomedical Engineering, University of Rochester, Rochester, NY; Department of Pathology & Laboratory Medicine, University of Rochester Medical Center, Rochester, NY; Department of Orthopaedics & Physical Performance, University of Rochester Medical Center, Rochester, NY

**Keywords:** Periostin, scar tissue, tendon, extracellular matrix

## Abstract

Tendon injuries are a major clinical problem, with poor patient outcomes caused by abundant scar tissue deposition during healing. Myofibroblasts play a critical role in the initial restoration of structural integrity after injury. However, persistent myofibroblast activity drives the transition to fibrotic scar tissue formation. As such, disrupting myofibroblast persistence is a key therapeutic target. While myofibroblasts are typically defined by the presence of αSMA+ stress fibers, αSMA is expressed in other cell types including the vasculature. As such, modulation of myofibroblast dynamics via disruption of αSMA expression is not a translationally tenable approach. Recent work has demonstrated that Periostin-lineage (Postn^Lin^) cells are a precursor for cardiac fibrosis-associated myofibroblasts. In contrast to this, here we show that Postn^Lin^ cells contribute to a transient αSMA+ myofibroblast population that is required for functional tendon healing, and that Periostin forms a supportive matrix niche that facilitates myofibroblast differentiation and persistence. Collectively, these data identify the Periostin matrix niche as a critical regulator of myofibroblast fate and persistence that could be targeted for therapeutic manipulation to facilitate regenerative tendon healing.

## Introduction

Tendons facilitate skeletal movement through force transfer from muscle to bones and are able to bear immense physiological loads due to their hierarchically organized collagen-rich matrix structure [1]. However, injuries to tendons are common: for example, one instance of tensile overloading may cause spontaneous Achilles tendon rupture [2-5], while flexor tendons of the hands are the tendons most frequently injured by acute trauma [6-8]. Independent of the mechanism of injury, tendons heal poorly, including a propensity for a scar-mediated, fibrotic healing response, rather than regeneration of the native tendon structure, which often prevents full functional restoration after injury [9, 10]. Moreover, the fibrovascular scar tissue response results in a healed tendon that is mechanically inferior to healthy tendon, which increases the incidence of re-rupture or repair failure. Despite this burden, the fibrotic healing process is not well-understood, which has led to a lack of consensus biological treatments to improve tendon healing.

Myofibroblasts are a key regulator of both physiological wound healing and tissue fibrosis. Myofibroblasts are capable of a high degree of extracellular matrix (ECM) production, while their inherent contractile ability modulates the stiffness of the wound matrix, thereby modulating the behavior of other cells embedded within this matrix [11-13]. Myofibroblasts are typically defined by *de novo* expression of αSMA in mechanosensitive stress fibers, which facilitates sensing and response to local ECM perturbations. While myofibroblasts are crucial for proper wound healing via ECM synthesis and remodeling, inappropriate accumulation and/or persistence of myofibroblasts at the site of injury is a major trigger for the switch to fibrotic progression[11, 14, 15]. Myofibroblasts can drive the release of latent TGF-β from the ECM, while TGF-β promotes myofibroblast differentiation, thereby creating a pro-fibrotic feedback loop [16-21]. In addition, physiological resolution of healing is inhibited by lack of myofibroblast clearance, either via apoptosis, or reversion to their basal fibroblastic state [22-25]. Thus, modulating the behavior of myofibroblasts is an attractive option to prevent fibrosis during tendon injury. However, manipulation of myofibroblast function via genetic mouse models to modulate αSMA expression is confounded by several factors. αSMA is expressed by other cell types, including smooth muscle cells in the vasculature [26]. Therefore, mouse models using an αSMA-Cre driver would cause off-target effects, and necessitates the identification of more translationally tractable markers of the myofibroblast fate. Recently, expression of the matricellular protein, Periostin (Postn) has been identified as a potential marker of activated myofibroblasts in cardiac fibrosis [27-30], with Periostin-lineage (Postn^Lin^) cells giving rise to myofibroblasts, while other studies have demonstrated that the Periostin matrix itself promotes and maintains myofibroblast differentiation [31, 32], and mediates tissue fibrosis [33-35]. Therefore, we sought to determine if Postn^Lin^ cells contributed to a similar population of pro-fibrotic myofibroblasts in the tendon, defined the role of the Periostin matrix niche in myofibroblast differentiation and persistence during tendon healing, and assessed the therapeutic potential of disrupting the formation of the Periostin matrix via inducible depletion of Postn^Lin^ cells.

## Methods

### Study Approval and Mouse Models

All studies were carried out in strict accordance with the recommendations in the Guide for the Care and Use of Laboratory Animals of the National Institutes of Health. All animal procedures were approved by the University Committee on Animal Research (UCAR) at the University of Rochester.

To trace Periostin-lineage (Postn^Lin^) cells, *Periostin*-Mer-iCre-Mer mice (Periostin^MCM^ #029645, Jackson Laboratories, Bar Harbor, ME) (kindly provided by Dr. Eric Small) were crossed to ROSA-Ai9 mice (#007909, Jackson Laboratories). The ROSA-Ai9 strain expresses the tdTomato fluorescent reporter following Cre-mediated deletion of a loxP-flanked STOP cassette. Following recombination, all Cre-expressing cells and their progeny permanently express tdTomato. To deplete Postn^Lin^ cells, *Periostin*^MCM^ mice were crossed to ROSA-DTA mice (#009669, Jackson Laboratories), which results in deletion of the floxed STOP cassette preceding the DTA transgene and subsequent expression of diphtheria toxin in all *Cre*-expressing cells. *Periostin*^MCM^; Rosa-Ai9^F/+^; Rosa-DTA^F/+^ mice were used for all experiments. The specific TMX (100mg/kg, i.p. injection) regimen used for each experiment is provided in each relevant figure.

### Murine model of tendon injury and repair

Male and female mice aged 10–12 weeks underwent complete transection and surgical repair of the flexor digitorum longus (FDL) tendon as previously described [36]. Briefly, mice were given a preoperative sub-Q injection of 15-20μg of sustained-release buprenorphine to manage pain throughout healing, and Ketamine (60mg/kg) and Xylazine (4mg/kg) given for anesthesia. Following removal of hair and sterilization of the surgical site, the flexor digitorum longus tendon was severed at the myotendinous junction (MTJ) to reduce strain on the repair site. The FDL in the hind paw was exposed via a skin-deep incision on the posterolateral surface and the FDL tendon was located and completely transected. The tendon was repaired using 8-0 suture with a modified Kessler pattern. The skin was then closed with a 5-0 suture. Mice were monitored and given analgesics post-operatively as needed.

### Histology and immunofluorescence

Following sacrifice, hind paws (n=3-5 per timepoint) were dissected just above the ankle, and skin carefully removed from both the plantar and dorsal surfaces. Hind paws were fixed in 10% formalin for 72 hours, decalcified in Webb-Jee EDTA for two weeks, then routinely processed and embedded in paraffin. Three-micron sagittal sections were cut, de-waxed, rehydrated, and probed with antibodies for tdTomato (1:500, AB8181, SICGEN), αSMA-CY3 (1:200, C6198, Sigma Life Sciences, St. Louis, MO, USA), αSMA-FITC (1:500, F3777, Sigma Life Sciences), HSP47 (1:250, ab109117, Abcam), or Periostin (1:200, ab14041, Abcam). The following secondary antibodies were used: Donkey anti-rabbit Rhodamine-Red-X (for Hsp47) (1:200, #711-296-152, Jackson Immuno), Donkey anti-goat Rhodamine-Red-X (for tdTomato) (1:200, #705-296-147, Jackson Immuno), Donkey anti-goat 488 (for tdTomato) (1:200, #705-546-147, Jackson Immuno), Donkey anti-rabbit 647 (for Periostin) (1:200, #711-606-152).

Nuclei were counterstained with NucBlue™ Live ReadyProbes™ Reagent (Hoechst 33342) (#R37605, ThermoFisher) and imaging was performed using a VS120 Virtual Slide Microscope (Olympus, Waltham, MA, USA). Select slides were also visualized using a Laser Scanning Confocal Microscope I (FV1000 Olympus, Center Valley, PA, USA) to examine the cellular interaction with the Periostin matrix. For the Masson’s Trichrome staining, coverslips were gently removed in PBS from slides previously stained and imaged for immunofluorescence, and then subsequently stained with Masson’s Trichrome to examine collagen organization.

### Measurement of gliding function

Following sacrifice, hindlimbs were harvested at the knee. The medial side of the hindlimb was carefully dissected to free the FDL from both the tarsal tunnel and myotendinous junction, and proximal end secured between two pieces of tape with cyanoacrylate. The distal tendon was loaded via the tape with a range of weights from 0g to 19g used to flex the metatarsophalangeal (MTP) joint, and the MTP flexion angle was recorded upon application of each weight. MTP Flexion Angle and Gliding Resistance were calculated as previously described, with lower MTP Flexion Angle and higher Gliding Resistance corresponding to impaired gliding function, consistent with increased scar tissue and peritendinous adhesion formation [37, 38].

### Quantification of biomechanical properties

FDL tendons were harvested from hind paws following gliding testing under a dissecting microscope. First the superficial flexor tendon was removed, the length of the FDL tendon including the repair site was carefully freed from surrounding connective tissues and cut just prior to the bifurcation into the digits. Discrepancy in sample number between measurement of gliding function and biomechanical testing resulted from tendon rupture during this detachment step, possibly due to strong adhesions to connective tissue. Each tendon fragment was prepared for uniaxial testing by placing two pieces of sandpaper on each end of the tendon and securing with cyanoacrylate (Superglue, LOCTITE). All the steps above were performed with the tissue periodically submerged in PBS to avoid any potential tissue drying. Each gripped tendon was transferred into a semi-customized uniaxial microtester (eXpert 4000 MicroTester, ADMET, Inc., Norwood MA). The microtester, along with the sample, was transferred to an inverted microscope (Olympus BX51, Olympus) to visualize the tendon and quantify the gauge length, width, and thickness. The gauge length of each sample was set as the end-to-end distance between opposing sandpaper edges and was the same for all samples tested. The cross-section of the tendon was assumed to be an ellipse, where the width of the tissue represents the major axis and the thickness of the tissue represents the minor axis. Based on the optically measured width and thickness of each tendon, the area of the elliptical cross-section was computed. A uniaxial displacement-controlled stretching of 1% strain per second until failure was applied. Load and grip-grip displacement data were recorded and converted to stress-strain data, and the failure mode was tracked for each mechanically tested sample. The load-displacement and stress-strain data were plotted and analyzed to determine structural (stiffness) and material (modulus) properties. Specifically, the slope of the linear region from the load displacement graph was determined to be the stiffness of the tested sample. The slope of the linear region from the stress-strain graph was taken to equal the elastic modulus parameter of each tested tendon. Note that this calculation assumes that stress and strain are uniform within each specimen.

### Western Blot Analysis

The tendon repair site was excised from the hind paw of *Periostin*-MCM; DTA F/+ mice at D14 following injury, along with 1–2 mm of native tendon on either side, then immediately snap-frozen in liquid nitrogen for storage at -80C until processing. Tissue (n=3 tendon per sample) was homogenized in RIPA buffer (including 1X protease/phosphatase inhibitor, Cell Signaling #58725) to extract the total protein with the Biomasher II system and Powermasher II electric stirrer (Nippi Inc, Tokyo Japan). Samples were spun down at 13000xg for 5 minutes, and supernatant transferred to a new tube to remove collagenous debris. A BCA protein assay was performed to quantify protein yield, following kit instructions (BCA Protein Assay kit, ThermoFisher #23225) and 20µg loaded into each well of a NuPAGE 4-12% Bis-Tris Gel (Invitrogen, #NP0335BOX, Carlsbad, CA). Gels were transferred to a membrane using the iBlot2 system (Invitrogen, IB23001), and then probed sequentially with anti-Periostin (1:1000, ab14041, Abcam) and the loading control β-Actin (1:2000, Cell Signaling, #4967) in 5% nonfat dry milk in TBST. Imaging was performed using the Pico: SuperSignal West Pico PLUS Chemiluminescent Substrate (Thermo Scientific, # 34580) and bands quantified with ImageJ.

### Primary tendon cell isolation and in vitro studies

C57BL/6J mice (#000664) mice were obtained from the Jackson Laboratory (Bar Harbor, ME, USA) and used for all primary tenocyte isolation protocols. For each experiment, FDL tendons (n=4) were aseptically excised, pooled, and collagenase digested (0.075% collagenase I; #C6885, Sigma) in fibroblast growth medium-2 (FGM-2; #CC-3132, Lonza, Basel, Switzerland) for one hour at 37°C with stirring. The collagenase mixture was then filtered at 70 μM and cells were pelleted, resuspended in FGM-2, and plated in collagen-coated plates (rat tail collagen type 1, 5 μg/cm^2^; #354236, Corning, Tewksbury, MA). Cells were maintained under standard culture conditions (37°C, 5% CO_2_, 90% humidity), with complete media exchanges every other day, and upon reaching 70% confluence, cells were passaged using 0.05% trypsin-EDTA (#25300–054, Gibco, Waltham, MA). All experiments were performed on cells at P1 or P2. To determine how Periostin may impact tenocyte differentiation to αSMA+ myofibroblasts, 4-well chamber slides (Nunc Lab-Tek II Chamber Slide System, Thermo Fisher) were pre-coated with collagen I alone as above, or with the addition of 5µg/mL recombinant Periostin (R&D Systems, #2955-F2) overnight at 4°C. Primary tenocytes were diluted to 10,000 cells/mL and 500uL was seeded in each well of a 4-chamber well slide (Nunc Lab-Tek II Chamber Slide System, Thermo Fisher) precoated with collagen alone, or with the addition of 5ug/mL recombinant Periostin (R&D Systems, #2955-F2). After 24h incubation, cells were fixed in 3% PFA in PBS for 10 minutes, washed, permeabilized with 0.1% Triton-X in PBS, then stained with α-SMA-Cy3 (1:200, C6198, Sigma Life Sciences) overnight at 4°C. The next day, nuclei were counterstained with NucBlue™ Live ReadyProbes™ Reagent (Hoechst 33342) (#R37605, ThermoFisher), and slides imaged with the VS120 Virtual Slide Microscope. Ten fields were chosen per well for point-counting of fully differentiated myofibroblasts (identified by α-SMA stress fibers) and quantification in ImageJ. The average of the 10 fields for each well constituted a single data point. Experiments were performed in biological triplicate and experimental duplicate.

### RNA Isolation and qPCR analysis

For in vitro studies, 200uL TRIzol was added to each well of the 24-well plate seeded with primary tenocytes and treated as described above. A P1000 pipette tip was used to scrape cells into solution for lysis, and suspension collected to a 1.5mL Eppendorf tube, followed immediately by RNA isolation or storage at -20°C. RNA isolation was carried out using the Direct-zol RNA Microprep Kit (Zymo Research #R2062) by column purification and including a DNAse step to remove DNA contamination. cDNA was generated with 500 ng of RNA using an iScript cDNA synthesis kit (BioRad, Hercules CA). Quantitative PCR was run with gene specific primers (Table 1), and data were normalized to β*-*actin (*Actb*) and expression in WT samples using the 2^-ΔΔCt^ method to determine the relative fold gene expression [39]. All experiments were done in biological triplicate and experimental duplicate.

**Table 1.**
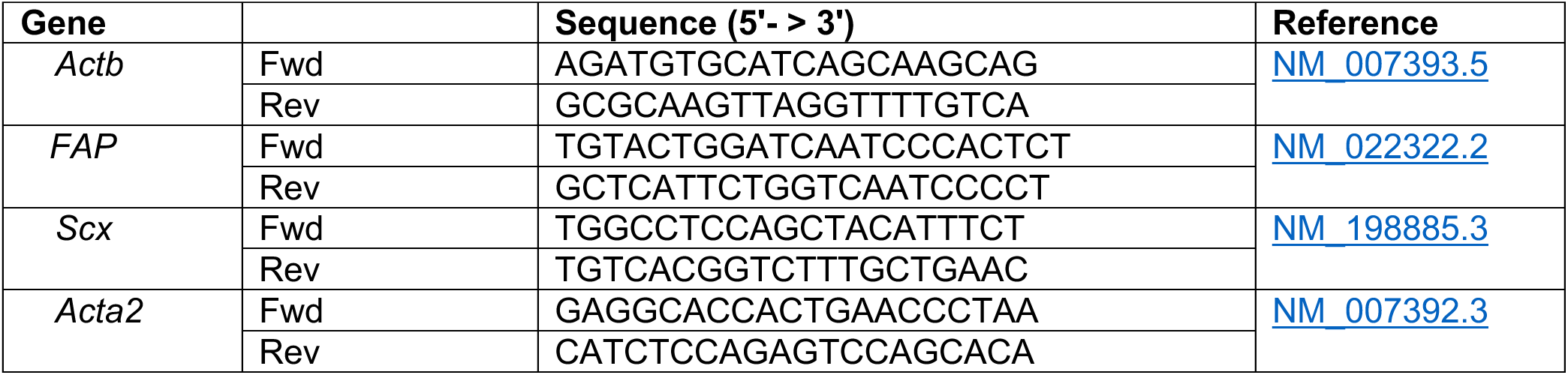
qPCR Primer Sequences.

### Human tissue samples

Human flexor tendon samples were isolated in accordance with our IRB approved protocol (RSRB #54231). Healthy control tendons (n=2) were harvested from the digits following traumatic hand amputations. Tendon scar tissue was isolated during ‘tenolysis’ surgery (n=3) to remove function-limiting peritendinous adhesions and scar tissue that formed following primary surgical repair of flexor tendon lacerations. Tenolysis procedures occurred between 3-12 months after the primary tendon repair surgery.

### Statistics

Statistically significant differences between genotypes or treatments in *in vitro* and *in vivo* studies were assessed by unequal variance un-paired t-test. All analyses were conducted using GraphPad Prism software (v9.2., La Jolla CA). Data are presented as mean ± SD, and p-values ≤ 0.05 were considered significant, with the following conventions: *=p ≤ 0.05, **=p ≤ 0.01, and ***=p ≤ 0.001. Mice were randomly allocated to specific experimental outcome metrics prior to surgery. Outlier data points for tested for using GraphPad Prism software using the ROUT method, and the Q value set at 1%.

## Results

### αSMA+ myofibroblasts reside within a Periostin matrix niche in mouse and human tendon scar tissue

We first defined the expression pattern of Periostin in healthy mouse (Figure 1A) and human (Figure 1B) flexor tendon. During homeostasis, minimal Periostin expression was observed in murine tendon, while Periostin was strongly expressed in the epitenon of human tendon. We then examined the Periostin expression profile during healing, and the relationship between Periostin and αSMA+ myofibroblasts (Figure 1C). During the early phase of healing (D3 and D7), Periostin expression was observed around the native tendon stubs, and very few αSMA+ myofibroblasts were present. By D10, robust expansion of the Periostin matrix occurred, concomitant with an increase in αSMA+ myofibroblasts. Interestingly, αSMA+ myofibroblasts were very often found embedded within the secreted Periostin matrix (Figure 1D) suggesting ongoing cell-matrix communication throughout healing. While the location and extent of Periostin matrix changed substantially from D10-28, this close proximity between the Periostin matrix and αSMA+ myofibroblasts was retained (Figure 1C). We then examined the relationship between Periostin and αSMA+ myofibroblasts in human tendon scar tissue. This peritendinous scar tissue, formed in response to an acute tendon injury and following primary repair of the tendon, restricts tendon function and is isolated during a ‘tenolysis’ or scar removal surgery. Within this tissue there were regions with robust Periostin expression, which overlapped with regions enriched for αSMA+ myofibroblasts (Figure 1E), further supporting a strong association between the Periostin-rich matrix and myofibroblasts.

**Figure 1.**
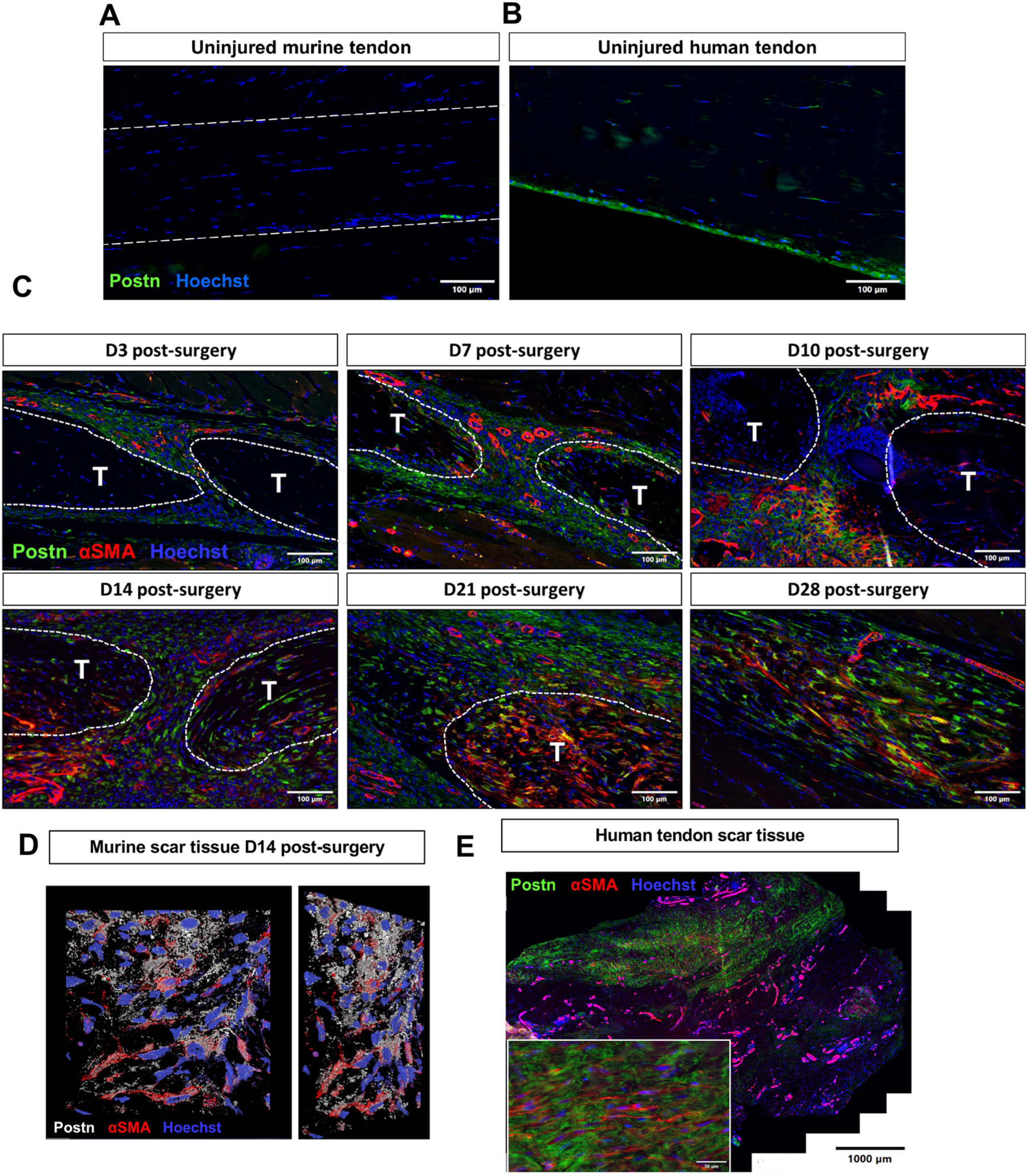
Periostin expression is dynamic during tendon healing. (A) During homeostasis minimal Periostin (Postn, green) expression is observed in C57Bl/6J murine flexor digitorum longus (FDL) tendons, while Periostin expression was observed in the epitenon of uninjured human flexor tendon samples (B). (C) C57Bl/6J mice underwent FDL repair surgery and were harvested for histology at D3, 7, 10, 14, 21, and 28. Tissue sections were stained for α-SMA (red), secreted Periostin (green), and Hoescht (nuclei, blue). Following injury, areas of Periostin expression correspond with localization of αSMA+ myofibroblasts. (D) A 2D representation of a 3D confocal z-stack imaged from an area of the murine scar tissue at D14 post-surgery. Periostin staining is shown in white, αSMA in red, and nucle in blue. (E) Human tenolysis tissue was processed for paraffin sectioning and stained for αSMA (green), Periostin (red), and Hoescht (nuclei, blue). Inset is magnified from the area enclosed by a box in the left overall image.

### Periostin-lineage cells give rise to a transient myofibroblast population during tendon healing

To determine whether Periostin-lineage (Postn^Lin^) cells give rise to persistent myofibroblasts, Periostin^MCM^+; Rosa-Ai9^F/+^ mice were initiated on tamoxifen chow at the time of tendon repair surgery to continuously label and trace Postn^Lin^ cells (Figure 2A). Postn^Lin^ cells were not observed in uninjured contralateral control tendons (Figure 2B). By D7, Postn^Lin^ cells expand along the epitenon region, and co-localization with αSMA was seen closer to the injured tendon end (Figure 2C). This pattern became more obvious by D14, with a population of Postn^Lin^; αSMA+ cells also observed within the bridging scar tissue. However, during the remodeling stage (D28), minimal αSMA expression was observed in the Postn^Lin^ cells surrounding the tendon ends and in the bridging tissue (Figure 2C), and Postn^Lin+^ cells were retained in the thickened epitenon throughout healing. Collectively, these data demonstrate that Postn^Lin^ cells give rise to a subpopulation of transient myofibroblasts during the proliferative phase of tendon healing, but do not contribute to the persistent myofibroblast population during late healing.

**Figure 2.**
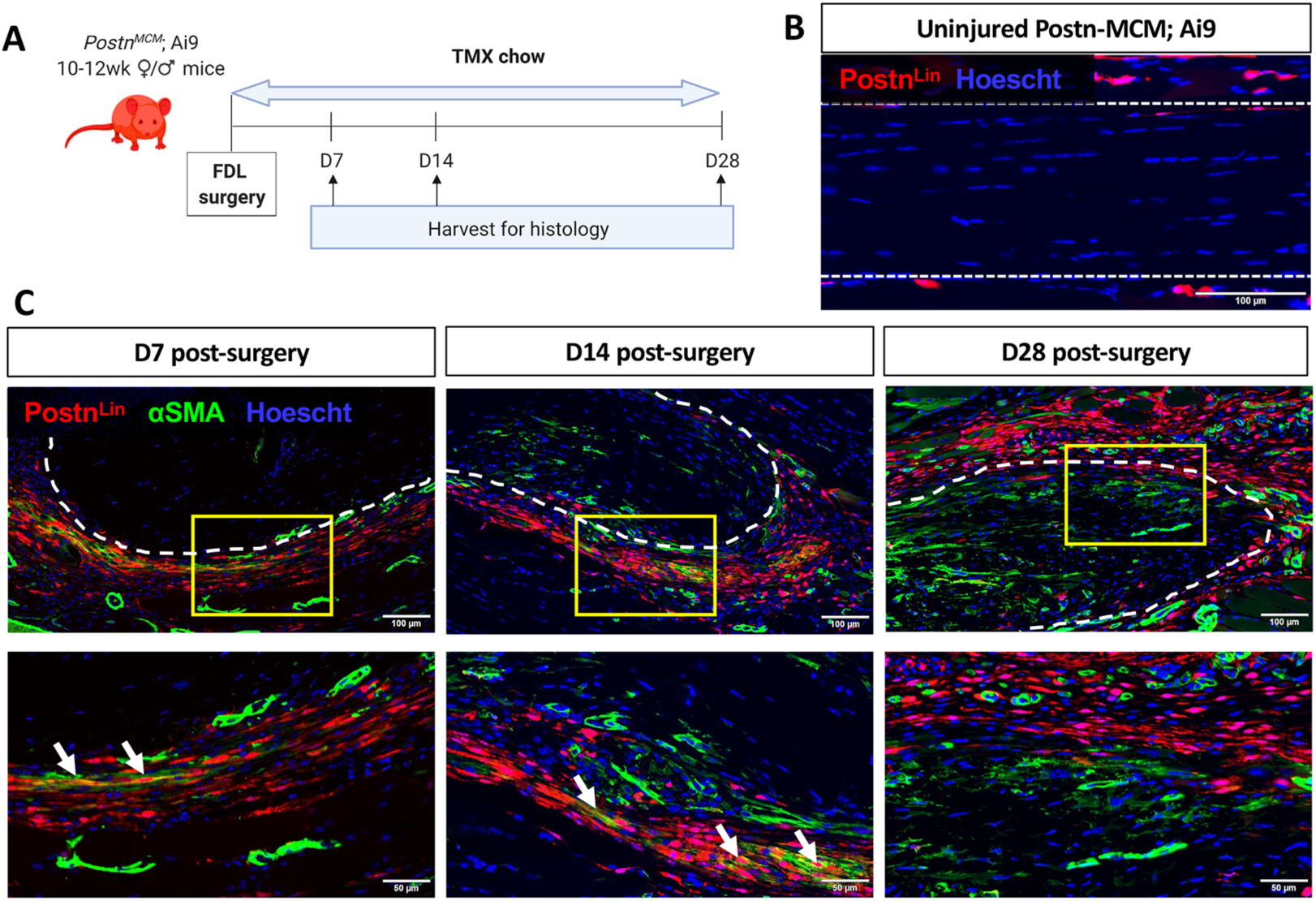
Postn^Lin^ cells transiently contribute to the αSMA+ myofibroblast pool during tendon healing. (A) To trace Periostin-lineage (Postn^Lin^) cells, Postn^MCM^; Rosa-Ai9 reporter mice were fed tamoxifen chow from the time of surgery until harvest at D28 post-surgery. (B) Very few Postn^Lin^ cells (red) were observed in uninjured contralateral control tendons. Nuclei are stained blue with Hoescht. (C) At D7 and D14 post-surgery, a small population of Postn^Lin^ cells (red) express αSMA (green), indicative of myofibroblast fate (white arrows), however, by D28 no overlap between Postn^Lin^ cells and αSMA+ myofibroblasts is observed. Native tendon stubs are outlined with dotted white lines.

### Postn^Lin^ cell activation peaks at approximately D10 post-injury and Postn^Lin^ cells give rise to spatially and functionally heterogeneous subpopulations

Given that Postn^Lin^ cells are not the primary contributor to the persistent myofibroblast population, and that persistent myofibroblasts are embedded in a robust Postn matrix in mouse and human tendon scar tissue, we sought to determine the temporal window(s) in which new Postn-expressing cells are added to the overall Postn^Lin^ pool to elaborate the Postn-rich matrix. Four cohorts of Postn^Lin^ mice underwent Tmx labelling during periods encompassing early to late healing (Figure 3A). Labelling from D3-4 resulted in very few Postn^Lin^ cells at D5. In contrast, labelling from D7-8 resulted in the identification of a robust Postn^Lin^ population (Figure 3B). The Postn^Lin^ population at D16, identified via labelling from D14-15, was slightly decreased relative to that labelled from D7-8, and a marked reduction in the Postn^Lin^ population was observed when these cells were labelled from D21-22 and assessed at D23 (Figure 3B). Collectively, these data suggest that the largest contribution to the overall Postn^Lin^ pool occurs from ∼D9-16.

**Figure 3.**
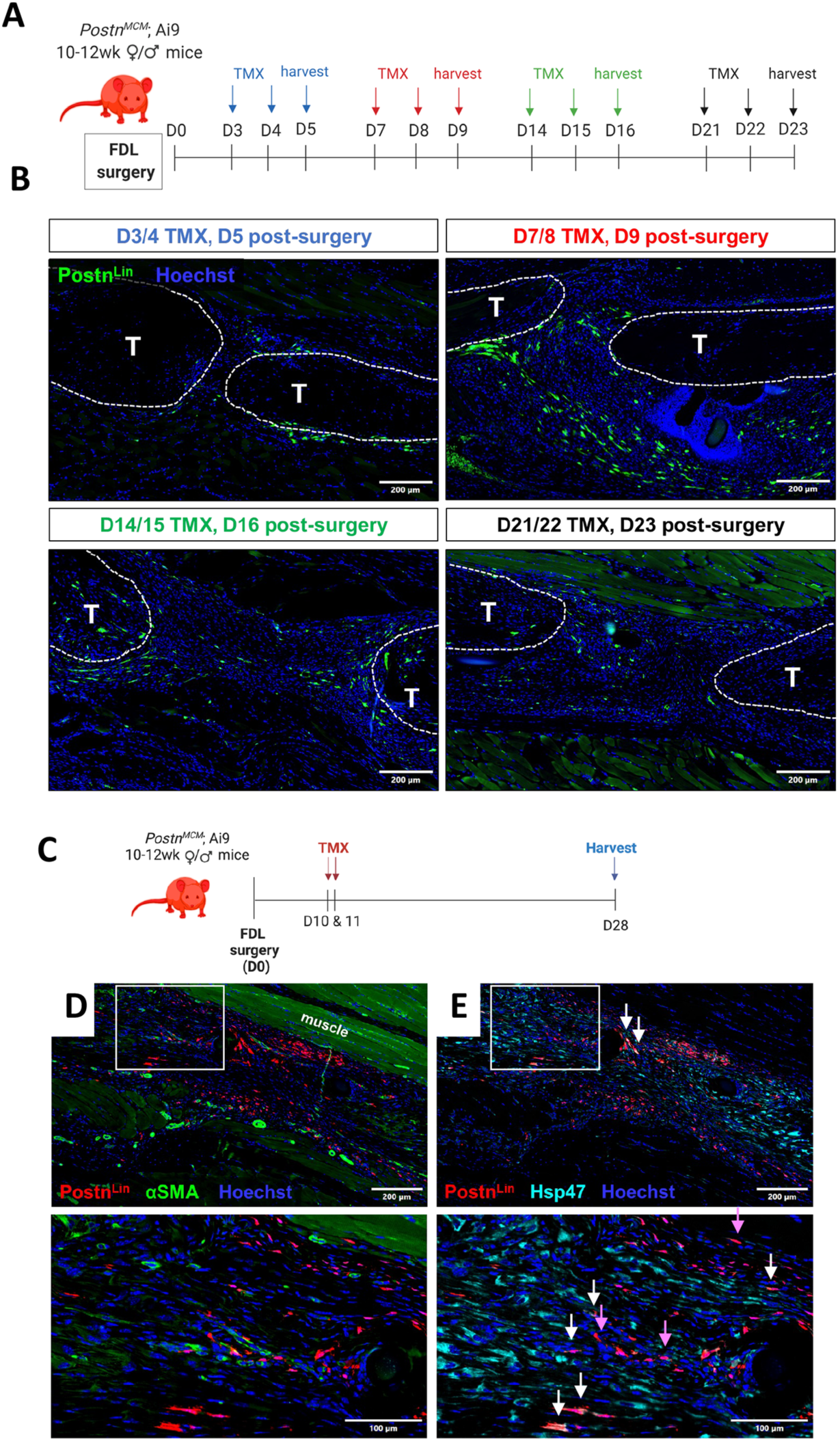
Postn^Lin^ cells are primarily added to the pool between D7-10, and make time-dependent contributions to healing. (A & B) Temporally-dependent contributions to the overall Postn^Lin^ pool were defined using four different Tmx treatment and harvest regimens. Tmx was given for two days and hind paws were harvested on the third day of the regimen for immunofluorescent analysis, with the following regimens used: TMX D2 & 3, harvest D4 (blue); TMX D7 & 8, harvest D9 (red); TMX D14 & 15, harvest D16 (green); TMX D21 & 22, harvest D23 (black). Postn^Lin^ cells are shown in green, αSMA+ myofibroblasts in red, and nuclei are stained blue with Hoescht. Tendon ends are outlined by dotted white lines. (C) Based on the large addition of Postn^Lin^ to the healing environment at this time, Postn^Lin^ cells were labelled via Tmx administration on D10 & 11, with tendons harvested at D28. (D) At D28, minimal overlap between this subset of Postn^Lin^ cells (red) and αSMA+ myofibroblasts (green) was observed, indicating this subset of Postn^Lin^ cells does not contribute to the persistent myofibroblast population. (E) Several Postn^Lin^ cells actively expressed Hsp47 at this time (white arrows), while there was also a large population of Postn^Lin^ cells that did not express Hsp47 (purple arrows).

To better understand the functional role of the Postn^Lin^ cells that are added to the overall Postn^Lin^ cells during this period, we labelled Postn^Lin^ cells at D10-11 post-surgery and then traced these cells until D28 (Figure 3C). Interestingly, cells labelled at D10-11 did not contribute to the thickened epitenon (Figure 3D, white box), as was observed with tamoxifen chow, suggesting that the Postn^Lin^ epitenon cells are labelled prior to D10 in the healing process. While Postn^Lin^ cells labelled at D10/11 persist to D28, consistent with the Tmx chow lineage trace, they do not give rise to the persistent αSMA+ myofibroblast population (Figure 3D). To examine whether this αSMA^-^ subset of Postn^Lin^ cells contribute to collagen production, we examined Hsp47 expression (collagen chaperone protein). Hsp47 was widespread throughout the tendon ends and scar tissue with Hsp47+ Postn^Lin^ cells in the bridging scar tissue (Figure 3E, white arrows). However, there was also a subset of Postn^Lin^ cells lining the bridging scar tissue that lack Hsp47 (Figure 3E, purple arrows). Collectively, these data suggests that Postn^Lin^ cells make temporally and spatially-distinct contributions to healing, including giving rise to a transient population of myofibroblasts, elaboration of both a myofibroblast-associated matrix niche, and a peripheral epitenon-associated matrix niche lacking myofibroblasts.

### Periostin matrix is sufficient to induce primary tenocyte activation and myofibroblast differentiation

Given the close association between the Periostin matrix and myofibroblasts, we assessed the ability of Periostin to promote tenocyte activation and myofibroblast differentiation in vitro. Expression of the fibroblast activation marker Fibroblast Activation Protein (*FAP*) was significantly increased (∼14-fold, p<0.01) in tenocytes cultured on rPeriostin-coated plates, without a corresponding increase in *Scx* expression (p=0.3), suggesting that this activation is occurring without any change in tenogenic potential. In addition, a significant increase in αSMA expression (2-fold, p=0.0009) was observed in tenocytes on rPeriostin-coated plates, relative to control. No change in Col3a1 expression was observed (p=0.49) (Figure 4A). Tenocytes cultured on rPeriostin-coated plates underwent a significantly greater degree of myofibroblast differentiation (αSMA stress fiber+ cells; +47% increase, p = 0.011), relative to tenocytes grown on collagen alone (Figure 4B). Collectively, these data suggest that Periostin may act as an extracellular niche to potentiate fibroblast activation and myofibroblast differentiation via cell-matrix crosstalk.

**Figure 4.**
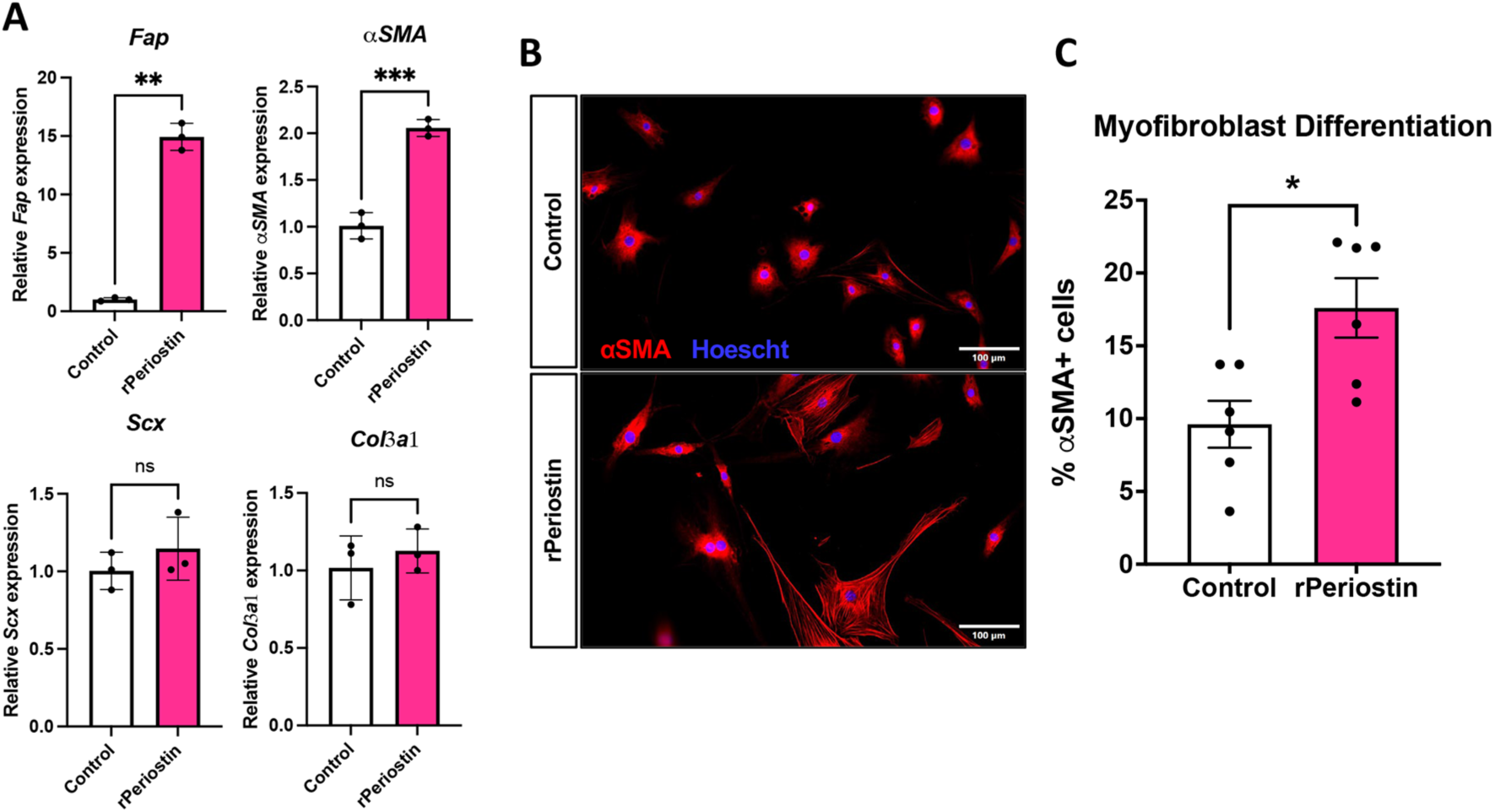
Periostin matrix drive tenocyte activation and differentiation *in vitro*. Primary murine tenocytes were seeded onto tissue culture plates coated with either Collagen I alone (control), or Collagen I with the addition of 10µg/mL recombinant Periostin (rPeriostin). (A) Changes in expression of genes associated with fibroblast activation (*Fap*), myofibroblast differentiation (*αSMA)*, tenogenesis (*Scx)*, and a healing-associated matrix (*Col3a1)* were assessed in tenocytes cultured on control and rPostn-coated plates. N=3 biological replicates (cells isolated from individual mice), with three technical replicates per biological sample. (B & C) The impact of rPeriostin on myofibroblast differentiation was assessed via immunostaining for αSMA (red), and nuclei were stained blue with Hoescht. The proportion of mature myofibroblasts (αSMA+ stress fibers) were quantified relative to the overall cell count.n=6 biological replicates, with three technical replicates per biological sample. (*) indicates p<0.05, (**) indicates p<0.01, (***) indicates p<0.001, (NS) indicates lack of statistical significance.

### Inhibition of Periostin matrix elaboration impairs tendon functional recovery

As we have shown that a Periostin matrix is sufficient to drive tenocyte-myofibroblast differentiation in vitro, we investigated the therapeutic potential of disrupting myofibroblast persistence via depletion of Postn^Lin^ cells and inhibition of subsequent Periostin matrix elaboration. We hypothesized that depletion just prior to the peak addition to the overall Postn^Lin^ pool (∼D9-16) would be most beneficial and cells were depleted from D7-10 (Figure 5A). However, this approach was not sufficient to disrupt the established Periostin matrix, and did not alter αSMA+ myofibroblast content (Figure 5B). Consistent with the persistence of the Periostin matrix and lack of impact on the myofibroblast environment, no changes in tendon gliding function (Figure 5C), or reacquisition of mechanical properties (Figure 5D) were observed with cell depletion from D7-10.

**Figure 5.**
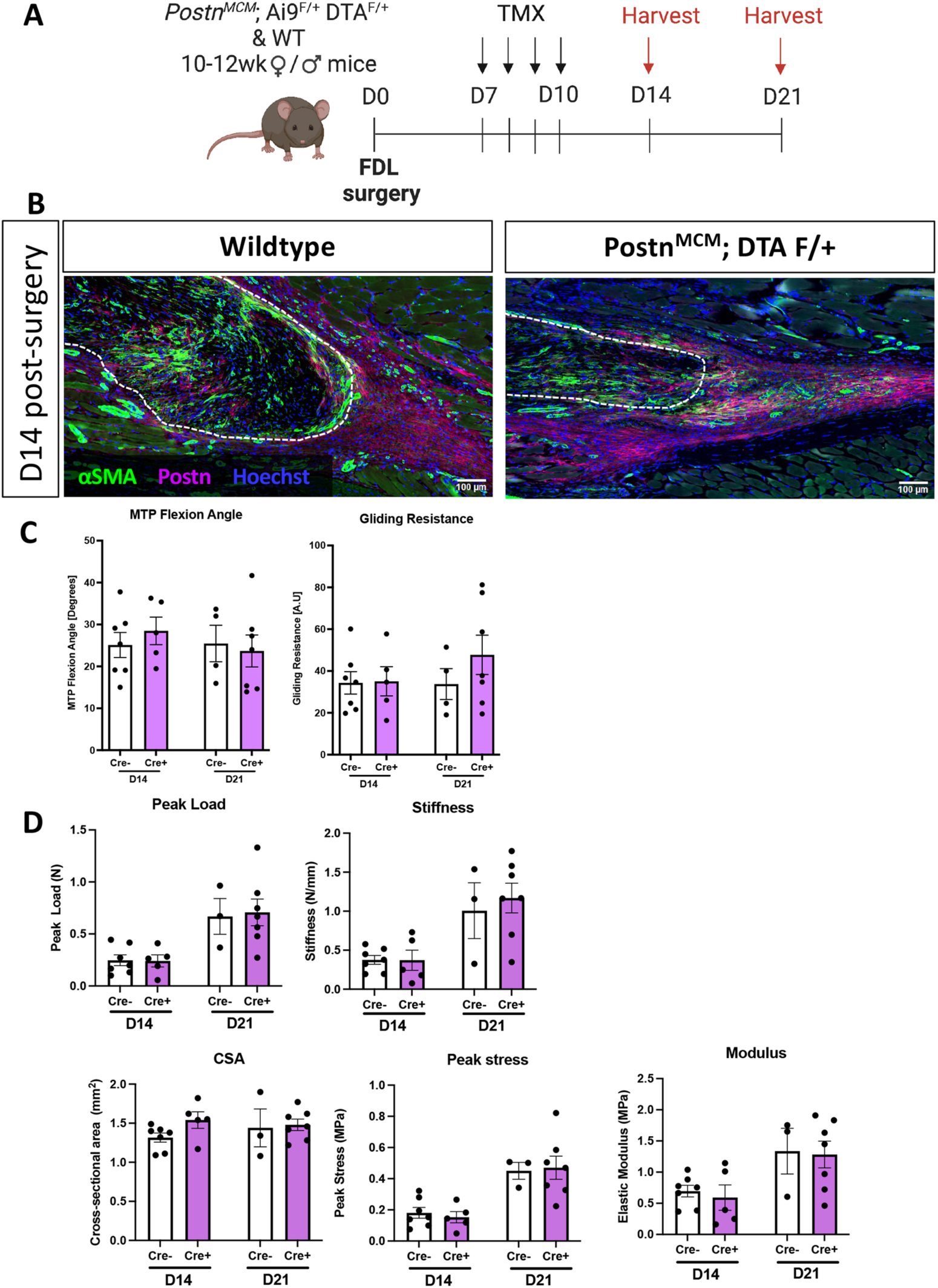
Depletion of Postn^Lin^ cells from D7-10 post-injury is insufficient to disrupt the mature Periostin matrix. (A) To deplete Postn^Lin^ cells and disrupt the established Periostin matrix, Postn-MCM; Ai9 F/+; DTA F/+ mice were given tamoxifen from D7-10 post-surgery, and tendons were harvested at D14 and 21. (B) Deletion of Postn^Lin^ cells from D7-10 did not disrupt the established Periostin matrix (purple), or alter myofibroblast content (αSMA+ cells, green). Nuclei are stained blue with Hoescht. (C) No changes in gliding function, including MTP Flexion Angle, or Gliding Resistance were observed between groups. (D) No changes in mechanical or material properties were observed between groups.

Given the lack of Periostin matrix disruption with the initial depletion regimen, we widened the depletion window to D5-14 (Figure 6A). This resulted in an 83% decrease in Periostin protein in the healing tendon as measured by western blot (Figure 6B). In addition, a significant 52% decrease in Periostin+ area in the scar tissue was observed in depleted tendons, relative to WT, via immunofluorescence analysis (p=0.023) (Figure 6C), further supporting the efficacy of this depletion regimen. A corresponding reduction in αSMA staining was also observed, and although not significant (-26%, p>0.05), this is consistent with impaired myofibroblast differentiation with decreased Periostin matrix and Postn^Lin^ cells (Figure 6D). However, linear regression analysis showed significant positive correlation between αSMA and Periostin staining (R^2^ = 0.6763, P = 0.023), further supporting the requirement of a Periostin matrix for αSMA staining indicative of myofibroblast differentiation (Figure 6E).

**Figure 6.**
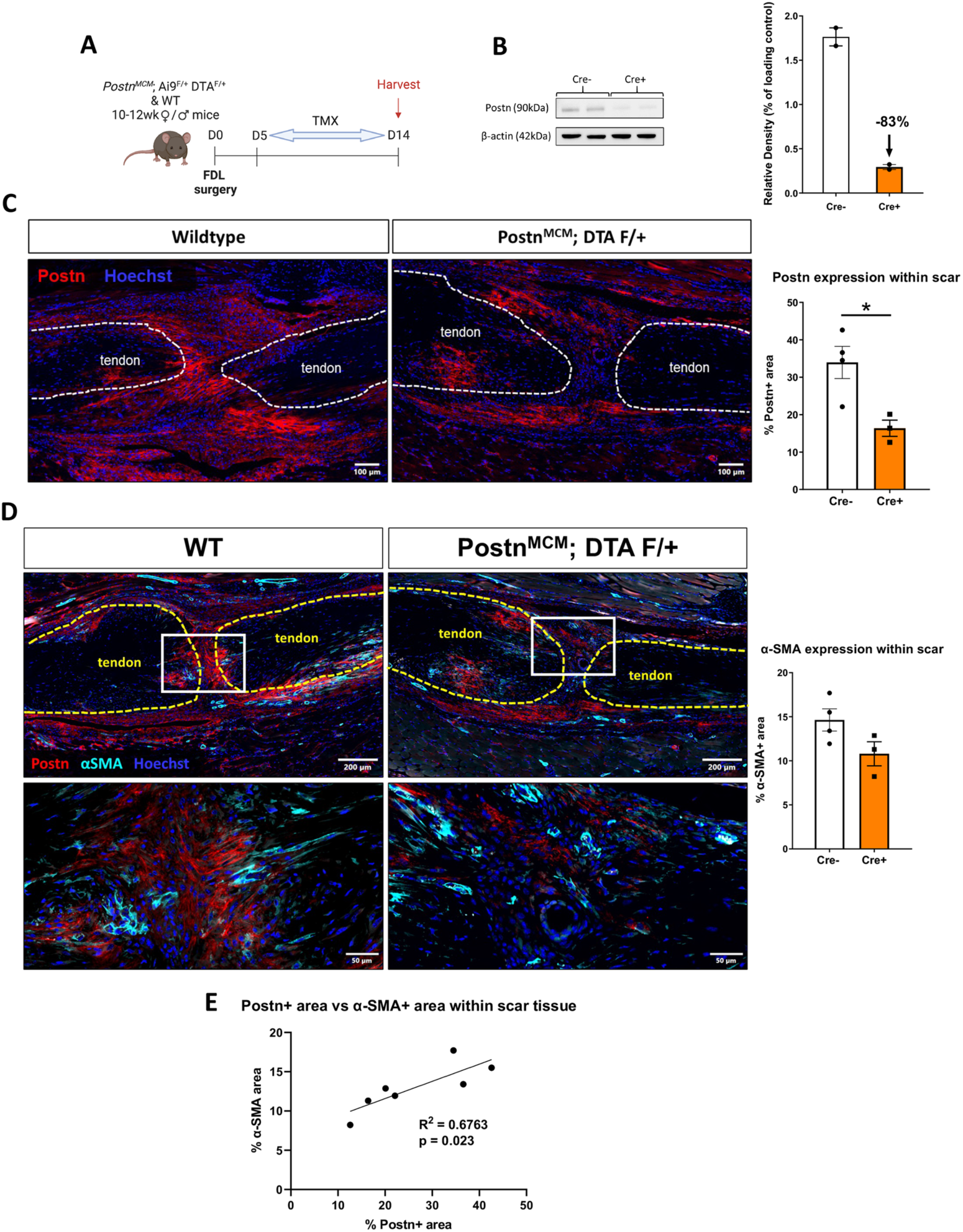
Depletion of Postn^Lin^ cells from D5-14 post-injury disrupts formation of the Periostin matrix. (A) In an attempt to disrupt the Periostin matrix, Postn-MCM; Ai9^F/+^; DTA^F/+^ mice were given tamoxifen from D5-14 post-surgery, and hind paws were harvested at D14. (B) Western blotting confirms an 83% decrease in Postn protein using this depletion regimen (n=2 biological replicates with a pooling of three tendons per biological replicate). (C & D) Wildtype control and Postn^Lin^-depleted tendons were stained for Periostin matrix (red), and αSMA (D, cyan). Nuclei are stained blue with Hoescht. A significant reduction in Periostin+ area was observed in depleted tendons, while a trending but not significant decline in αSMA was also observed. Less αSMA and Periostin staining is qualitatively observed in the depleted mice. (E) Linear regression analysis of αSMA and Periostin staining showed a significant positive correlation between between myofibroblast content and Periostin matrix abundance.

Functionally, Postn^Lin^ cell depletion from D5-14 resulted in a marked impairment of gliding function (Gliding Resistance: +47%, p = 0.0044, MTP Flexion Angle: -34%, p = 0.0041) as well as peak load and stress (Peak Load: -50%, p = 0.0471 and Peak Stress: -50%, p = 0.0493) at D14 (Figure 7A & B). Collectively, these data suggest that maintenance of a Periostin matrix during the proliferative phase of healing is necessary for functional tendon recovery after injury. Consistent with functional impairments, Postn^Lin^ cell depletion resulted in substantial disruption in tissue morphology. More specifically, integration of scar tissue at the muscle-tendon interface was observed (Figure 7C, yellow arrows), compared to a clearer delineation in wildtype repairs (Figure 7C, white arrows).

**Figure 7.**
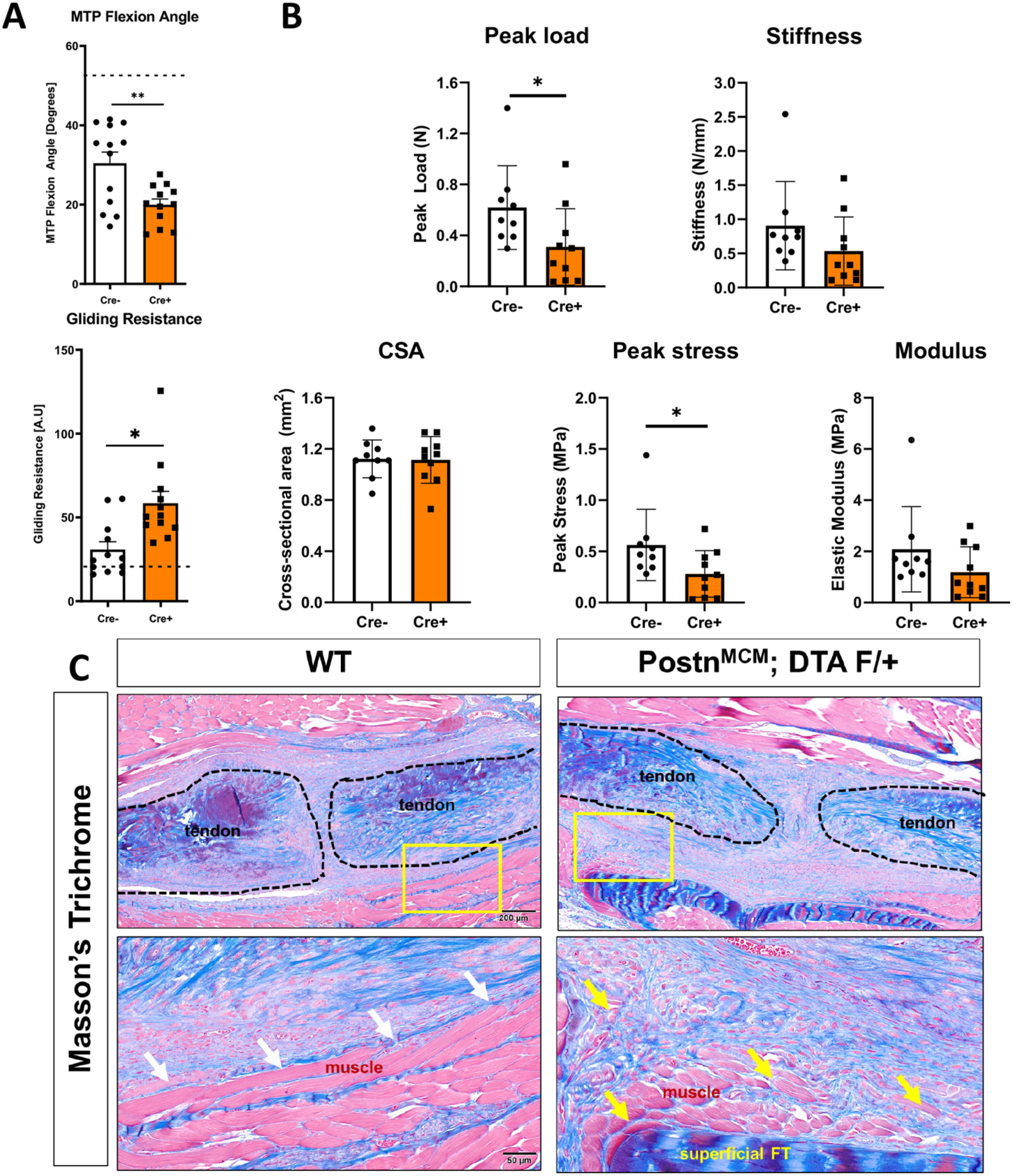
Depletion of Postn^Lin^ cells from D5-14 post-injury impairs functional and biomechanical recovery during tendon healing. (A) Postn^Lin^ cell depletion from D5-14 resulted in substantial declines in tendon range of motion as shown by a significant decrease in MTP joint flexion angle and a significant increase in Gliding Resistance, relative to wildtype repairs (n=12 per genotype). (B) Assessment of recovery of mechanical and material properties demonstrated significant declines in peak load at failure and peak stress, relative to wildtype. No changes were observed in stiffness, cross-sectional area (CSA) or elastic modulus, relative to wildtype. (n=9-10 per genotype). (C) Histologically, intercalation of the tendon scar tissue into the muscle at the injury site was consistently observed in depleted mice (yellow arrows), in comparison to the clearly delineated border in wildtype littermates (white arrows), as shown by Masson’s Trichrome staining.

## Discussion

Myofibroblasts are critical regulators of the wound healing process, including in the tendon, due to their ability to synthesize, contract, and remodel the provisional matrix that restores some structural integrity to the injured tissue. Conversely, persistent myofibroblast activity is associated with the transition to a fibrotic healing response. As such, modulation of the myofibroblast environment is an attractive target to facilitate regenerative healing in many tissues. However, because myofibroblasts are typically defined by the presence of αSMA+ stress fibers, and αSMA is broadly expressed in multiple cell types including the vasculature, manipulating myofibroblasts by altering αSMA expression confounds data interpretation and is not translationally tenable.

Therefore, identification of additional markers of myofibroblasts, and an increased understanding of the tissue-specific mechanisms that control myofibroblast differentiation and persistence are needed to identify feasible strategies to enhance healing. Recent work has demonstrated that Postn^Lin^ cells give rise to myofibroblasts following myocardial infarction [40], and genetic deletion of Periostin blunts myofibroblast differentiation in multiple cells and tissue types [41, 42]. Therefore, we examined the role of both Postn^Lin^ cells and the Periostin matrix during tendon healing. Here we show that while Postn^Lin^ cells are the not the predominant contributors to the persistent myofibroblast population, a robust Periostin matrix forms in response to tendon injury in both mice and humans, the Periostin matrix acts as a niche for myofibroblasts, and inhibiting formation of the Periostin matrix, via Postn^Lin^ cell depletion, disrupts myofibroblast differentiation and impairs tendon healing.

While the role of Periostin in tendon development and homeostasis is not entirely clear, recent work has demonstrated that global deletion of Periostin disrupts tendon homeostasis, including impairments in collagen fibrillogenesis and acquisition of mechanical properties [43], while overexpression of Periostin in mesenchymal cells results in modest formation of tendon-like tissue in an ectopic implantation model [44]. During healing, increased Periostin expression is observed after injury in both Achilles tendons [45] and the Medial Collateral Ligament [46], consistent our data demonstrating the development of Periostin-rich areas in healing murine tendons and clinical isolates of peritendinous scar tissue. Moreover, Periostin^-/-^ results in delayed and mechanically inferior Achilles tendon healing, relative to wildtype littermates [43], consistent with delayed healing in both palatal tissue [41] and skin [42]. However, it remains to be determined whether these tendon healing deficits are primarily due to homeostatic disruptions in Periostin^-/-^ tendons, or the lack of Periostin in the healing process. To partially address this, we depleted Postn^Lin^ cells at different stages of the healing process, which results in an inducible decrease in Periostin and avoids homeostatic disruption prior to injury. Importantly, these data demonstrate time-dependent effects of Postn^Lin^ cell depletion. While Postn^Lin^ depletion from 7-10 days post-surgery is insufficient to disrupt the established Periostin matrix that is deposited prior to day 7, depletion of this subset of Postn^Lin^ cells does not impair the healing process suggesting that Postn^Lin^ cells themselves are not required for functional healing during this period. In contrast, depletion of Postn^Lin^ cells from days 5-14 post-surgery is sufficient to disrupt elaboration of the Periostin matrix, resulting in impaired tendon healing and establishing the requirement for both the initial formation of the Periostin matrix as well as Postn^Lin^ cells during this window. Consistent with this, recent work has demonstrated the efficacy of a recombinant Periostin-laden engineered construct to enhance tendon healing, likely due to enhanced Tendon Stem Progenitor Cell (TSPC) migration [47].

Despite the requirement for Postn^Lin^ cells and initial Periostin matrix elaboration in tendon healing, and the encouraging results with rPeriostin-laden constructs to further improve tendon healing [47], the strong association between persistent myofibroblasts and the Periostin matrix in clinical tendon scar tissue samples supports the need for additional investigation to determine the potential therapeutic benefit of targeting the persistent Periostin matrix as an anti-fibrotic therapy, and highlights the important consideration of therapeutic timing. Global deletion of Periostin reduces fibroblast, myofibroblast, and immune cell recruitment during palatal healing [41], and reduces myofibroblast content during skin healing [42]. However, this approach disrupts progress of the entire wound healing response, and does not facilitate interrogation of the role of Periostin once the healing response has switched from physiological to fibrotic. While technical challenges limited our ability to disrupt the Periostin matrix once it had formed, the lack of impact on healing of depleting Postn^Lin^ cells from D7-10 potentially supports the feasibility of targeting the Periostin matrix once it has been elaborated. Again, timing is likely to be critical as Periostin is required for collagen maturation and cross-linking [30] and regulation of collagen fibrillogenesis [48-50], however disruption of the persistent myofibroblast niche may represent an as yet explored area for facilitating myofibroblast clearance or reversion and promoting restoration of a physiological, tenogenic program.

The difference in sustained myofibroblast fate of Postn^Lin^ cells between this study and prior work is likely due to differences in timing of both analysis and labelling of the Postn^Lin^ population. Prior work traced Postn^Lin^ cells following two weeks of tamoxifen chow, while mice in this study remained on tamoxifen chow for the entire experiment to more comprehensively label and trace the broad Postn^Lin^ pool. The contribution of non-Postn^Lin^ cells to the myofibroblast pool is consistent with contributions from a heterogeneous pool of cells contributing to the myofibroblast fate in multiple tissues [51-53]. While it has long been recognized that fibroblast heterogeneity is an active determinant in pathologies ranging from fibrotic wound healing to cancer[54-59], myofibroblast heterogeneity remains relatively understudied, despite long-term evidence [60]. However, the advent of single-cell and spatial RNA sequencing techniques has given researchers the tools to better understand and appreciate the high degree of cellular heterogeneity in various disease processes [61-63] and is an important focus for future tendon research.

One of the more surprising findings of this study was the presence of Postn^Lin^ cells in the thickened epitenon after injury. Recent work [64] has demonstrated the bi-fated potential of epitenon cells, including a tenogenic progenitor population and a pro-fibrotic population that appears on the periphery of the injury site. Continuously labeled Postn^Lin^ cells demonstrates robust presence in the thickened epitenon and along the margins of the bridging tissue by D28 post-injury. Intriguingly, while myofibroblasts are typically implicated as the primary pro-fibrotic population, their absence from the epitenon-derived exterior capsule population, and the concomitant enrichment for Periostin matrix and Postn^Lin^ cells in this area suggests a previously unappreciated, non-myofibroblast role for Postn^Lin^ cells in peritendinous adhesion formation.

One of central limitations of this study is that the experimental approach we used to disrupt the Periostin-matrix during healing concomitantly ablated Postn^Lin^ cells. In doing so, the cell environment and subsequent cell-cell communication patterns are altered. Future studies to disrupt the Periostin matrix, while retaining the Postn^Lin^ cell population are needed to clearly delineate between matrix and cell-autonomous functions of Periostin. Recent work has demonstrated the potential of cathepsin K to disrupt the Periostin matrix during fracture healing [65]. As such, future work is needed to assess the efficacy of this approach to modulate the mature Periostin matrix during tendon healing.

Collectively, these data demonstrate that Postn^Lin^ cells play a unique role during tendon healing, relative to other tissues. In healing tendons, Postn^Lin^ cells give rise to a transient population of αSMA+ myofibroblasts, rather than persistent population that is observed during cardiac fibrosis. Moreover, elaboration of a mature Periostin matrix serves as a critical niche to regulate myofibroblast differentiation and persistence during healing, and we have demonstrated the exterior epitenon-derived tissue associated with peritendinous adhesion formation is enriched for Periostin matrix and Postn^Lin^ cells. As such, we have established modulation of the Periostin matrix niche as a key regulator of fibrotic tendon healing and manipulation of this matrix as important therapeutic target to enhance the tendon healing process.

## Acknowledgements

This work was supported in part by NIH/ NIAMS F31 AR077398 (to JEA), K99 AR080757 (AECN), R01AR073169 and R01AR077527 (to AEL). The HBMI and BBMTI Cores were supported by NIH/ NIAMS P30 AR069655. The content is solely the responsibility of the authors and does not necessarily represent the official views of the National Institutes of Health.

## References

1. Eisner, L.E., et al., The Role of the Non-Collagenous Extracellular Matrix in Tendon and Ligament Mechanical Behavior: A Review. J Biomech Eng, 2022. 144(5).

2. Abdul Alim, M., et al., Achilles tendon rupture healing is enhanced by intermittent pneumatic compression upregulating collagen type I synthesis. Knee Surg Sports Traumatol Arthrosc, 2018. 26(7): p. 2021–2029.

3. Ahmad, J., M. Repka, and S.M. Raikin, Treatment of myotendinous Achilles ruptures. Foot Ankle Int, 2013. 34(8): p. 1074–8.

4. Barfod, K.W., Achilles tendon rupture; assessment of nonoperative treatment. Dan Med J, 2014. 61(4): p. B4837.

5. Hess, G.W., Achilles tendon rupture: a review of etiology, population, anatomy, risk factors, and injury prevention. Foot Ankle Spec, 2010. 3(1): p. 29–32.

6. Mehrzad, R., et al., The Economic Impact of Flexor Tendon Lacerations of the Hand in the United States. Ann Plast Surg, 2019. 83(4): p. 419–423.

7. Schöffl, V., A. Heid, and T. Küpper, Tendon injuries of the hand. World journal of orthopedics, 2012. 3(6): p. 62–69.

8. Hoxie, S.C., et al., The economic impact of electric saw injuries to the hand. J Hand Surg Am, 2009. 34(5): p. 886–9.

9. A Clinical, Biological, and Biomaterials Perspective into Tendon Injuries and Regeneration. Tissue Engineering Part B: Reviews, 2017. 23(1): p. 44–58.

10. Chartier, C., et al., Tendon: Principles of Healing and Repair. Semin Plast Surg, 2021. 35(3): p. 211–215.

11. Desmoulière, A., C. Chaponnier, and G. Gabbiani, Tissue repair, contraction, and the myofibroblast. Wound Repair Regen, 2005. 13(1): p. 7–12.

12. McAnulty, R.J., Fibroblasts and myofibroblasts: their source, function and role in disease. Int J Biochem Cell Biol, 2007. 39(4): p. 666–71.

13. Midwood, K.S., L.V. Williams, and J.E. Schwarzbauer, Tissue repair and the dynamics of the extracellular matrix. Int J Biochem Cell Biol, 2004. 36(6): p. 1031–7.

14. Gibb, A.A., M.P. Lazaropoulos, and J.W. Elrod, Myofibroblasts and fibrosis: mitochondrial and metabolic control of cellular differentiation. Circulation research, 2020. 127(3): p. 427–447.

15. Van De Water, L., S. Varney, and J.J. Tomasek, Mechanoregulation of the myofibroblast in wound contraction, scarring, and fibrosis: opportunities for new therapeutic intervention. Advances in wound care, 2013. 2(4): p. 122–141.

16. Braga, T.T., J.S. Agudelo, and N.O. Camara, Macrophages During the Fibrotic Process: M2 as Friend and Foe. Front Immunol, 2015. 6: p. 602.

17. Hou, J., et al., M2 macrophages promote myofibroblast differentiation of LR-MSCs and are associated with pulmonary fibrogenesis. Cell Commun Signal, 2018. 16(1): p. 89.

18. Yu, Y., et al., Exosomes From M2 Macrophage Promote Peritendinous Fibrosis Posterior Tendon Injury via the MiR-15b-5p/FGF-1/7/9 Pathway by Delivery of circRNA-Ep400. Front Cell Dev Biol, 2021. 9: p. 595911.

19. Hinz, B., It has to be the alphav: myofibroblast integrins activate latent TGF-beta1. Nat Med, 2013. 19(12): p. 1567–8.

20. Hinz, B., Tissue stiffness, latent TGF-beta1 activation, and mechanical signal transduction: implications for the pathogenesis and treatment of fibrosis. Curr Rheumatol Rep, 2009. 11(2): p. 120–6.

21. Rifkin, D.B., Latent transforming growth factor-β (TGF-β) binding proteins: orchestrators of TGF-β availability. Journal of Biological Chemistry, 2005. 280(9): p. 7409–7412.

22. Best, K.T., et al., NF-kappaB activation persists into the remodeling phase of tendon healing and promotes myofibroblast survival. Sci Signal, 2020. 13(658).

23. Hecker, L., et al., Reversible differentiation of myofibroblasts by MyoD. Experimental Cell Research, 2011. 317(13): p. 1914–1921.

24. Hinz, B. and D. Lagares, Evasion of apoptosis by myofibroblasts: a hallmark of fibrotic diseases. Nature Reviews Rheumatology, 2020. 16(1): p. 11–31.

25. Kisseleva, T., et al., Myofibroblasts revert to an inactive phenotype during regression of liver fibrosis. Proc Natl Acad Sci U S A, 2012. 109(24): p. 9448–53.

26. Zhao, W., et al., alpha-smooth muscle actin is not a marker of fibrogenic cell activity in skeletal muscle fibrosis. PLoS One, 2018. 13(1): p. e0191031.

27. Fu, X., et al., Specialized fibroblast differentiated states underlie scar formation in the infarcted mouse heart. The Journal of Clinical Investigation, 2018. 128(5): p. 2127–2143.

28. Kanisicak, O., et al., Genetic lineage tracing defines myofibroblast origin and function in the injured heart. Nature Communications, 2016. 7(1): p. 12260.

29. Khalil, H., et al., Fibroblast-specific TGF-beta-Smad2/3 signaling underlies cardiac fibrosis. J Clin Invest, 2017. 127(10): p. 3770–3783.

30. Schwanekamp, J.A., et al., Deletion of Periostin Protects Against Atherosclerosis in Mice by Altering Inflammation and Extracellular Matrix Remodeling. Arteriosclerosis, Thrombosis, and Vascular Biology, 2016. 36(1): p. 60–68.

31. Crawford, J., et al., Periostin induces fibroblast proliferation and myofibroblast persistence in hypertrophic scarring. Experimental dermatology, 2015. 24(2): p. 120–126.

32. Nikoloudaki, G., et al., Periostin and matrix stiffness combine to regulate myofibroblast differentiation and fibronectin synthesis during palatal healing. Matrix Biology, 2020. 94: p. 31–56.

33. O’Dwyer, D.N. and B.B. Moore, The role of periostin in lung fibrosis and airway remodeling. Cell Mol Life Sci, 2017. 74(23): p. 4305–4314.

34. Naik, P.K., et al., Periostin promotes fibrosis and predicts progression in patients with idiopathic pulmonary fibrosis. Am J Physiol Lung Cell Mol Physiol, 2012. 303(12): p. L1046–56.

35. Ashley, S.L., et al., Periostin regulates fibrocyte function to promote myofibroblast differentiation and lung fibrosis. Mucosal Immunol, 2017. 10(2): p. 341–351.

36. Ackerman, J.E. and A.E. Loiselle, Murine Flexor Tendon Injury and Repair Surgery. J Vis Exp, 2016(115).

37. Hasslund, S., et al., Adhesions in a murine flexor tendon graft model: Autograft versus allograft reconstruction. J Orthop Res, 2008. 26(6): p. 824–33.

38. Loiselle, A.E., et al., Remodeling of murine intrasynovial tendon adhesions following injury: MMP and neotendon gene expression. J Orthop Res, 2009. 27(6): p. 833–40.

39. Livak, K.J. and T.D. Schmittgen, Analysis of relative gene expression data using real-time quantitative PCR and the 2(-Delta Delta C(T)) Method. Methods, 2001. 25(4): p. 402–8.

40. Snider, P., et al., Origin of cardiac fibroblasts and the role of periostin. Circ Res, 2009. 105(10): p. 934–47.

41. Nikoloudaki, G., et al., Periostin and matrix stiffness combine to regulate myofibroblast differentiation and fibronectin synthesis during palatal healing. Matrix Biol, 2020. 94: p. 31–56.

42. Elliott, C.G., et al., Periostin modulates myofibroblast differentiation during full-thickness cutaneous wound repair. J Cell Sci, 2012. 125(Pt 1): p. 121–32.

43. Rolnick, K.I., et al., Periostin modulates extracellular matrix behavior in tendons. Matrix Biol Plus, 2022. 16: p. 100124.

44. Noack, S., et al., Periostin Secreted by Mesenchymal Stem Cells Supports Tendon Formation in an Ectopic Mouse Model. Stem Cells and Development, 2014. 23(16): p. 1844–1857.

45. Chamberlain, C.S., et al., Temporal healing in rat achilles tendon: ultrasound correlations. Ann Biomed Eng, 2013. 41(3): p. 477–87.

46. Chamberlain, C.S., et al., Gene profiling of the rat medial collateral ligament during early healing using microarray analysis. J Appl Physiol (1985), 2011. 111(2): p. 552-65.

47. Wang, Y., et al., Functional regeneration and repair of tendons using biomimetic scaffolds loaded with recombinant periostin. Nature Communications, 2021. 12(1): p. 1293.

48. Kudo, A., Periostin in fibrillogenesis for tissue regeneration: periostin actions inside and outside the cell. Cellular and molecular life sciences, 2011. 68(19): p. 3201–3207.

49. Norris, R.A., et al., Periostin regulates collagen fibrillogenesis and the biomechanical properties of connective tissues. J Cell Biochem, 2007. 101(3): p. 695–711.

50. Walker, J.T., et al., Periostin as a multifunctional modulator of the wound healing response. Cell and Tissue Research, 2016. 365(3): p. 453–465.

51. Hinz, B., et al., The myofibroblast: one function, multiple origins. Am J Pathol, 2007. 170(6): p. 1807–16.

52. Kuppe, C., et al., Decoding myofibroblast origins in human kidney fibrosis. Nature, 2021. 589(7841): p. 281-286.

53. Mack, M. and M. Yanagita, Origin of myofibroblasts and cellular events triggering fibrosis. Kidney international, 2015. 87(2): p. 297–307.

54. Costa, A., et al., Fibroblast heterogeneity and immunosuppressive environment in human breast cancer. Cancer cell, 2018. 33(3): p. 463–479. e10.

55. Fries, K.M., et al., Evidence of fibroblast heterogeneity and the role of fibroblast subpopulations in fibrosis. Clinical immunology and immunopathology, 1994. 72(3): p. 283–292.

56. Jelaska, A., D. Strehlow, and J.H. Korn. Fibroblast heterogeneity in physiological conditions and fibrotic disease. In Springer seminars in immunopathology. 2000. Springer.

57. LeBleu, V.S. and E.G. Neilson, Origin and functional heterogeneity of fibroblasts. The FASEB Journal, 2020. 34(3): p. 3519–3536.

58. Lynch, M.D. and F.M. Watt, Fibroblast heterogeneity: implications for human disease. J Clin Invest, 2018. 128(1): p. 26–35.

59. Mascharak, S. and M.T. Longaker, Fibroblast heterogeneity in wound healing: hurdles to clinical translation. Trends in molecular medicine, 2020. 26(12): p. 1101–1106.

60. Skalli, O., et al., Myofibroblasts from diverse pathologic settings are heterogeneous in their content of actin isoforms and intermediate filament proteins. Lab Invest, 1989. 60(2): p. 275–85.

61. De Micheli, A.J., et al., Single-cell transcriptomic analysis identifies extensive heterogeneity in the cellular composition of mouse Achilles tendons. Am J Physiol Cell Physiol, 2020. 319(5): p. C885–C894.

62. Deng, C.C., et al., Single-cell RNA-seq reveals fibroblast heterogeneity and increased mesenchymal fibroblasts in human fibrotic skin diseases. Nat Commun, 2021. 12(1): p. 3709.

63. Guerrero-Juarez, C.F., et al., Single-cell analysis reveals fibroblast heterogeneity and myeloid-derived adipocyte progenitors in murine skin wounds. Nature communications, 2019. 10(1): p. 1–17.

65. Nichols, A.E.C., et al., Epitenon-derived cells comprise a distinct progenitor population that contributes to both tendon fibrosis and regeneration following acute injury. bioRxiv, 2023.

65. Bonnet, N., et al., Cathepsin K Controls Cortical Bone Formation by Degrading Periostin. Journal of Bone and Mineral Research, 2017. 32(7): p. 1432–1441.

